# Streamlined genomes, not horizontal gene transfer, mark bacterial transitions to unfamiliar environments

**DOI:** 10.1101/2024.12.27.630308

**Authors:** Swastik Mishra, Martin J. Lercher

**Affiliations:** Institute for Computer Science and Department of Biology, Heinrich Heine University, Düsseldorf, Germany

**Keywords:** Horizontal Gene Transfer, Bacterial Evolution, Ecosystem Transitions, Pathogenicity Gains, Purifying Selection

## Abstract

Bacterial colonization of unfamiliar environments constitutes a drastic evolutionary transition, likely altering the strength and direction of selection. This process is often assumed to increase rates of horizontal gene transfer (HGT), a major driver of bacterial adaptation. Larger genomes are thought to facilitate such adaptations by providing broader functional repertoires and more integration sites for foreign DNA. Here, we systematically test these ideas across a broad bacterial phylogeny linked to environmental transitions inferred from metagenomic data. Contrary to expectation, we find that bacteria entering new environments typically have smaller genomes and experience lower rates of HGT. The reduction in HGT is fully explained by genome size, with no residual effect of environmental transitions once size is controlled for. These findings suggest that successful bacterial colonizers rely less on genomic plasticity through HGT than previously assumed, highlighting gaps in our understanding of microbial evolutionary dynamics.

## Introduction

Bacterial strains frequently adapt to new ecosystems, with prominent examples reported from gut microbiota (Zheng et al., 2020), skin microbiota (Lieberman, 2022), bacterial-fungal associations (Richter et al., 2024), and colonization of marine plastics (Caruso, 2020). Here, we explore the genomic correlates of such colonization events. Specifically, we test three expectations: (1) Because ecosystem transitions are often accompanied by population bottlenecks, we expect a temporary weakening of purifying selection at these events; Moreover, as outlined below, there are reasons to hypothesize that (2) colonization is associated with large genomes, and (3) is accompanied by bursts of horizontal gene transfers (HGT). Large genomes may facilitate transitions to new ecosystems by providing a broader “toolbox” of molecular functions that can be employed in new combinations to address novel challenges (Maslov et al., 2009). In contrast, bacteria with small genomes often exhibit niche specialization, which can limit their adaptability when environmental conditions shift beyond their optimized range (Serra Moncadas et al., 2024). In addition, larger genomes are more likely to contain homologous genes that can serve as landing pads for horizontal gene transfer (HGT), potentially enhancing access to foreign DNA (Taylor et al., 2024).

HGT plays a central role in bacterial adaptation by enabling the acquisition of genomic fragments – entire genes or operons – from other organisms (Pang and Lercher, 2019, Pál et al., 2005, Arnold et al., 2021). This mechanism allows for the rapid gain of novel traits and may be particularly important during ecological transitions. Environmental change has therefore been hypothesized to increase HGT rates (Engelstädter and Moradigaravand, 2014), not only by creating new selective pressures but also by exposing bacteria to unfamiliar pangenomes that can be accessed via HGT (Dmitrijeva et al., 2024).

Several studies have shown that environmental change can indeed trigger significant spikes in HGT rates (Woods et al., 2020, Goh et al., 2024, Dadeh Amirfard et al., 2024, Acar Kirit et al., 2020). However, these findings primarily derive from laboratory experiments focused on specific gene families, particularly antibiotic resistance genes, despite HGT’s known capacity to transfer virtually any bacterial gene (van Dijk et al., 2020, Coluzzi et al., 2023). Expanding this research to the entire pangenome in natural settings has proved challenging, as tracking the complete mobilome within microbial communities presents significant technical hurdles (Brito, 2021).

In this study, we tested these three expectations using the GOLD database (Mukherjee et al., 2023), which provides metagenomic data for hundreds of bacterial genomes across diverse environments. From these data, we inferred ecosystem gains and linked them to nucleotide changes, ancestral genome sizes, and HGT rates across a deep bacterial phylogeny. The data indicate that (i) purifying selection weakens during these transitions, as expected. Contrary to expectations, however, we found that (ii) genomes involved in transitions to new ecosystems tend to be small, and (iii) there is no evidence for increased HGT rates at such ecosystem transitions.

## Results

For the analyses below, we used extant bacterial genomic data obtained from the EggNOG database v6 (Hernández-Plaza et al., 2023), which clusters genes into non-supervised orthologous groups or *NOGs*, referred to as gene families below. For the inference of ecosystem gains, we used the GOLD database, which contains metagenomic data from a variety of ecosystems. We limited our analyses to bacterial genomes where enough information is available such that they have at least 3 ‘Ecosystem Type’ labels as defined in GOLD. Our dataset encompasses 80 diverse Ecosystem types, which range from natural to laboratory ecosystems. Examples of the 80 Ecosystem Types include ‘fermented vegetables’, ‘fetus’, ‘fish products’, ‘freshwater’, and ‘industrial wastewater’.

Our final dataset comprises 8197 gene families across 159 taxa, such that each gene family is present in at least 4 taxa. To build a genome (or species) tree, we retrieved gene family trees and sequence alignments from EggNOG. We inferred a genome tree from gene family trees of 233 single-copy gene families present in at least 95% of all taxa, using the summary-based species tree method ASTRAL-Pro 2 (Zhang and Mirarab, 2022). The branch lengths of this tree were estimated with IQ-TREE 2 (Minh et al., 2020) based on a concatenation of the multiple sequence alignments.

To map ecosystem gains on branches of the genome tree, we used the asymmetric Wagner parsimony algorithm implemented in Count (Csűös, 2010). While originally developed for inferring gene gains and losses from presence–absence patterns, this method is equally applicable to other discrete traits. We applied it to a presence–absence matrix of GOLD Ecosystem Type labels across genomes, allowing us to identify branches with ecosystem gains (EG) and those with no ecosystem gains (NEG).

### Purifying selection weakens during transitions to new ecosystems, but only for a subset of gene families

At transitions to new ecosystems, we expect increased rates of adaptation (positive selection) for a subset of environment-related genes and, simultaneously, decreased purifying selection due to population bottlenecks. Both of these changes in natural selection should affect nucleotide substitutions in protein-coding genes, increasing the ratio of non-synonymous to synonymous nucleotide substitutions, *ω* = *dN/dS. ω* is a widely used metric to assess the type and strength of selection that acts on protein-coding genes. *ω <* 1 indicates purifying selection, while *ω* = 1 indicates purely neutral evolution. *ω >* 1 would indicate strong positive selection. Even when a gene product is under strong positive selection, this selection typically affects only a small subset of amino acids, such as those near an enzyme’s active site, while most others remain subject to purifying selection. For this reason, the average *ω* value for a gene rarely reaches or exceeds 1.

To test the prediction of increased *ω* at ecosystem gains, we obtained nucleotide sequences for our dataset from NCBI. We used the RELAX algorithm from the HyPhy software package (Wertheim et al., 2015) to compare *ω* values between branches with and without ecosystem gains. RELAX is designed to assess whether the strength of natural selection on a gene has been relaxed or intensified along a specified set of “test” branches compared to “reference” branches. In our case, we defined the test branches as those with ecosystem gains (EG) and the reference branches as those without ecosystem gains (NEG). RELAX then fits a codon model to the entire phylogeny, introducing a selection intensity parameter, *k*, which modifies the selection pressure on the test branches relative to the reference branches. This parameter is an exponent of *ω*. If *k >* 1, selection is intensified on the test branches; if *k <* 1, selection is relaxed. A likelihood ratio test is used to compare models with and without this parameter (corresponding to *k* = 1).

For 3228 gene families, we could reliably map EggNOG taxon IDs to corresponding genomes across all relevant taxa using the NCBI Datasets API (Sayers et al., 2022) and successfully ran RELAX. As expected from a dominance of purifying selection, only a small percentage of amino acid sites show signs of positive selection (*ω >* 1), with EG branches having a significantly higher percentage of such sites (median 2.40%) compared to NEG branches (median 1.63%, two-sided Mann-Whitney *U* = 5709953.5, *p* = 9.71 × 10^−12^, *N* = 3228).

RELAX detected a statistically significant change in the intensity of selection in ecosystem gains for 8.15% (263 out of 3228) of gene families (adjusted p < 0.05 after correcting for multiple tests (Benjamini and Hochberg, 1995)). Of these, 62.36% (164 out of 263) showed a decrease in the strength of selection, while the rest indicated an intensification of selection (Figure 1). Thus, while the majority of gene families show no significant shift in selection intensity, approximately two-thirds of the minority that does exhibit changes that are consistent with either positive selection or a relaxation of purifying selection, as expected.

**Fig. 1.**
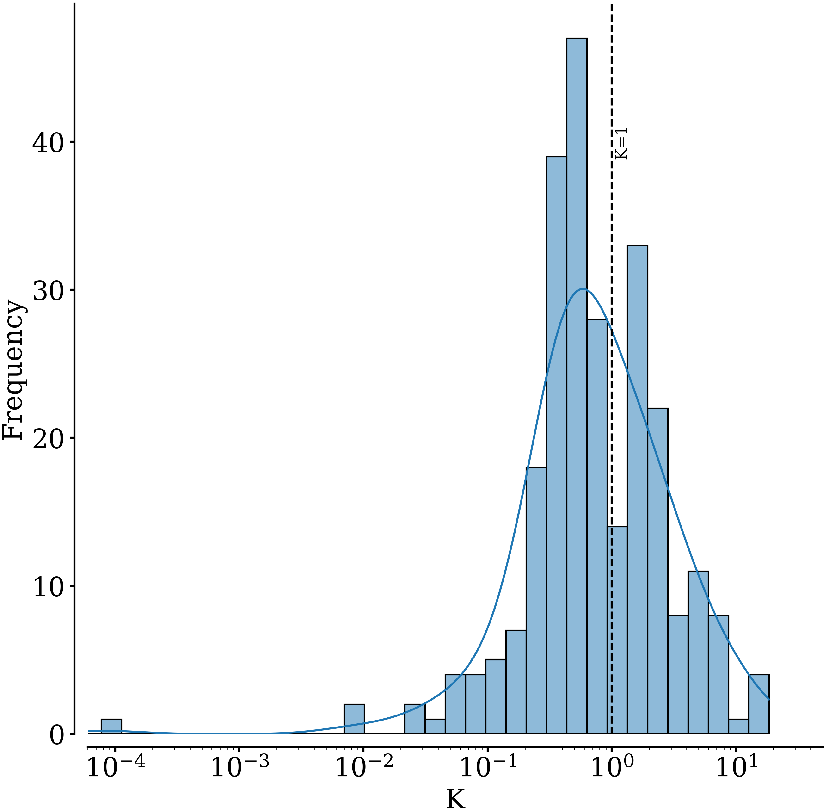
Distribution of *k* values for gene families with significant changes in selection intensity. Here, *k* indicates the change in strength of selection at ecosystem gains, with *k <* 1 indicating a relaxation of selection and *k >* 1 indicating an intensification of selection. The distribution is skewed towards values less than 1, indicating that the majority of gene families show a decrease in the strength of selection during ecosystem gains. The median *k* value is 0.61. The p-values for assessing significant change in *k* were corrected for multiple testing using the Benjamini-Hochberg method (Benjamini and Hochberg, 1995). ALT TEXT: Histogram of *k* values for gene families with significant changes in selection intensity at ecosystem gains.

### Bacteria transitioning to new ecosystems have smaller genomes and lower HGT rates

We next used the complete set of 8197 gene families to test our second prediction: that genomes undergoing transitions to new ecosystems tend to be larger than those that do not. Contrary to our expectation, branches with ecosystem gains tend to have smaller genomes, with a median of 3,800 genes compared to 5,395 in branches without such gains (Fig. 2, top marginal plot; two-sided Mann-Whitney *U* = 5884, *p* = 3.9 × 10^−12^, *N*_*EG*_ = 208, *N*_*NEG*_ = 108, respectively).

**Fig. 2.**
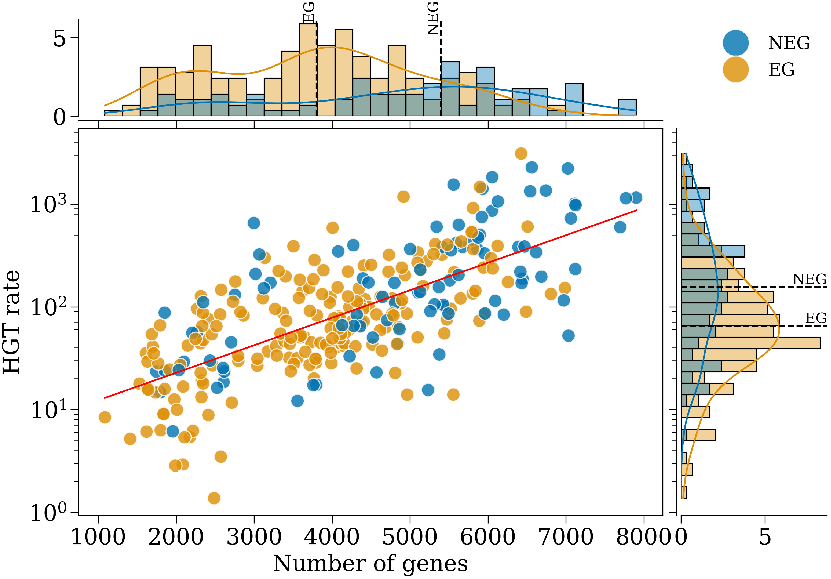
Bacteria transitioning to new ecosystems tend to have smaller genomes and lower HGT rates. As seen in the top marginal histogram, branches with ecosystem gains (EG, orange) tend to have a smaller genome size than branches with no ecosystem gains (NEG, blue). Branches with ecosystem gains also tend to have lower HGT rates, as shown in the right marginal histogram. Dotted lines show medians. Each point in the scatter plot represents a branch in the genome tree on which at least one gene was gained through HGT. Histograms show percentages. The regression line is *log*(*y*) = 6.65 + 6.16 *×* 10^−4^ · *x*, with a Pearson correlation coefficient *r* = 0.87. ALT TEXT: HGT rate is a function of genome size, as shown here in the scatter plot. Marginal plots show the distribution of genome sizes (top) and HGT rates (right) for branches with and without ecosystem gains.

Our third expectation was an increase in HGT rates at ecosystem gains. To robustly infer HGT events, we again used the complete gene set of 8197 gene families, complemented by the corresponding gene trees. A benchmark on empirical data demonstrated that Count’s maximum parsimony approach outperforms alternative HGT inference methods, including explicit, maximum-likelihood-based phylogenetic methods (Mishra and Lercher, 2024). We estimated the *HGT rate* for each branch of the genome phylogeny by dividing the number of inferred HGT events by the branch length. Accordingly, HGT rates are expressed as the number of HGT events per point substitution per amino acid site.

Figure 2 visualizes the HGT rates of branches with and without ecosystem gains, plotted against genome sizes. Contrary to the expectation that HGT rates would increase during transitions to new ecosystems, branches with ecosystem gains exhibited lower HGT rates than branches without (Figure 2, right marginal plot; two-sided Mann-Whitney *U* = 8306.5, *p* = 0.049, *N*_*EG*_ = 208, *N*_*NEG*_ = 108). This trend persists when the analysis is restricted to single-copy gene families (Supplementary Figure S1).

### Transitions to new ecosystems do not affect HGT rates when controlling for genome size

Thus, contrary to our expectations, we found that genomes undergoing transitions to new ecosystems tend to be small and experience low rates of HGTs. Given that smaller genomes generally exhibit lower rates of horizontal gene transfer (Fig. 2, scatter plot; Spearman *ρ* = 0.692, *p* = 1.4 × 10^−42^; see also Cordero and Hogeweg (2009)), it is conceivable that the lower HGT rates at ecosystem gains are simply a consequence of the smaller genome sizes. To test this conjecture, we fitted two linear models: a full model with HGT rate as a function of genome size and ecosystem gain, and a reduced model including genome size only. A likelihood ratio test comparing these nested models (*LRT* = 1.62, *p* = 0.203, *df* = 1) showed no significant improvement of the full model, indicating that ecosystem gain does not significantly affect HGT rate after accounting for genome size. Note that this analysis considered all gene families together. It is conceivable that ecosystem changes influence the HGT rate for specific functional categories; this is explored below.

### HGT rates do not change at transitions to pathogenic lifestyles

To test for the robustness of the finding that transitions to new ecosystems are not associated with changes in HGT rates, we used a similar pipeline for the examination of an independent dataset related to a different type of ecological change: the transition of *Escherichia coli* and *Shigella* strains from a commensal to a pathogenic lifestyle. Bacterial pathogenicity has been associated with changes in environmental conditions, such as a transition of a non-host associated organism to a host environment, but also transitions between different ecosystems within the host (Nuss et al., 2016, Chin et al., 2018, Brown et al., 2006).

To analyze transitions to pathogenicity, we examined 151 *E. coli* and *Shigella* genomes retrieved from NCBI, comprising 98 commensal and 53 pathogenic strains. For these genomes, we identified 5,503 orthologous gene families with OrthoFinder (Emms and Kelly, 2019) and estimated HGT rates using Count. Based on binary labels indicating whether a given genome is pathogenic, we also used Count to infer the gain and loss of pathogenicity. Pathogenicity gains were detected in just 14 of 300 evolutionary branches, a notably low frequency compared to ecosystem gains.

Mirroring trends observed in the ecosystem analyses, HGT rates increased with genome size (Spearman *ρ* = 0.442, *p* = 4.682 × 10^−14^). However, branches with pathogenicity gains did not have significantly smaller genomes (median 4,419 genes) than those without pathogenicity gains (median 4352 genes; two-sided Mann-Whitney *U* = 2344.5, *p* = 0.292, *N*_*P G*_ = 14, *N*_*NP G*_ = 287; Figure 3, top marginal histogram). Thus, whatever difference exists in HGT rates between branches with and without pathogenicity gains, they are unlikely to be caused by genome size effects.

**Fig. 3.**
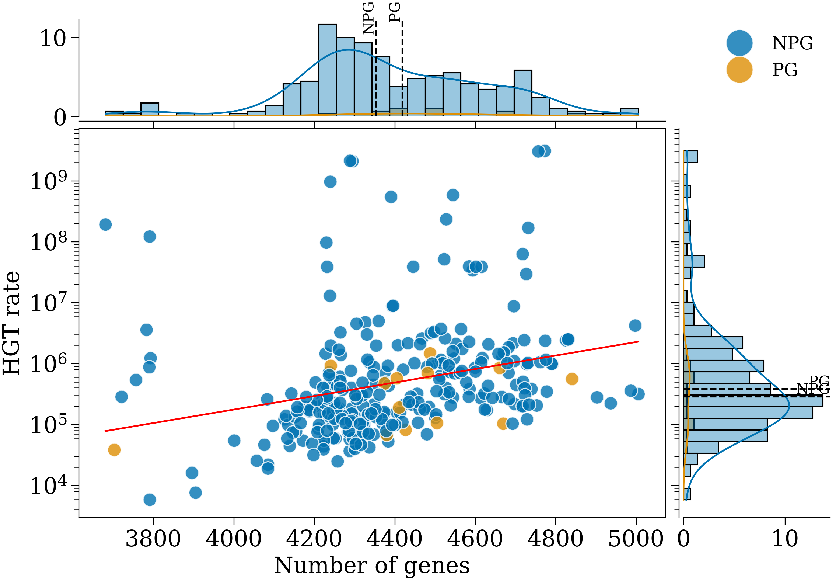
Transitions to pathogenic lifestyles are not associated with increased HGT rates or smaller genomes. As seen in the top marginal histogram, branches with pathogenicity gains (PG, orange) do not have significantly smaller genomes than branches without pathogenicity gains (NPG, blue). Branches with pathogenicity gains also do not have significantly higher HGT rates, as shown in the right marginal histogram. Dotted lines show medians. Each point in the scatter plot represents a branch in the genome tree on which at least one gene was gained through HGT. Histograms show percentages. The regression line is *log*(*y*) = 6.51 + 2.55 *×* 10^−3^ · *x*, with a Pearson correlation coefficient *r* = 0.1 and Spearman *ρ* = 0.442. ALT TEXT: HGT rate is a function of genome size, as shown here in the scatter plot. Marginal plots show the distribution of genome sizes (top) and HGT rates (right) for branches with and without pathogenicity gains.

While median HGT rates are marginally higher in branches with pathogenicity gains (3.819 × 10^−5^) than in those without (2.864 × 10^−5^), this difference is not statistically significant (two-sided Mann-Whitney *U* = 1835.0, *p* = 0.679, *N*_*P G*_ = 14, *N*_*NP G*_ = 287). In sum, ecological shifts – whether through adaptations to pathogenic lifestyles or the colonization of new ecosystems – are not associated with elevated HGT rates.

### Transitions to new ecosystems are associated with different HGT rates in only two functional categories of genes

When all gene families are analyzed collectively, ecosystem gains have no significant effect on HGT rates after controlling for genome size. However, ecosystem gains may still influence HGT rates in specific functional categories. To investigate this possibility, we examined each functional category defined in the Clusters of Orthologous Groups (COG) database (Tatusov et al., 2000, Galperin et al., 2021) that included at least 20 gene families in our dataset. We performed likelihood ratio tests (LRTs) as described earlier, comparing a model that includes both genome size and ecosystem changes as linear predictors of HGT rates with a nested model that considers only genome size.

Only the COG categories ‘Amino acid transport and metabolism’ (adjusted *p* = 3.1 × 10^−5^ after correction for multiple testing (Benjamini and Hochberg, 1995)) and ‘Transcription’ (adjusted *p* = 0.046) are significantly affected by ecosystem gains (Figure 4, Table S1). In both cases, HGT rates are lower on branches with ecosystem gains.

**Fig. 4.**
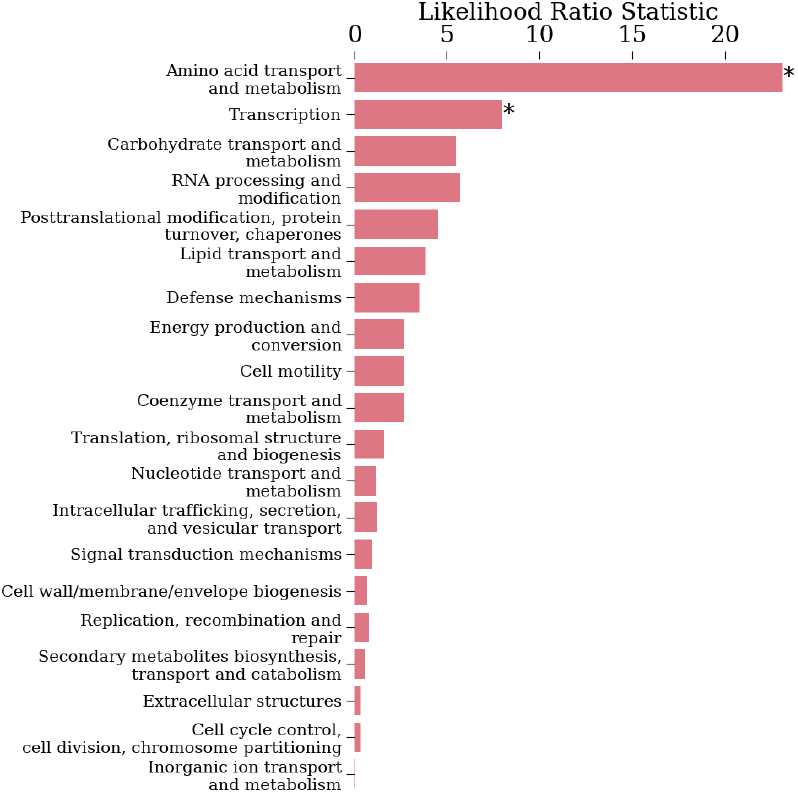
Functional categories with significant ecosystem gain effects on HGT rates. This barplot displays Log Likelihood Ratio statistics from likelihood ratio tests (LRTs) comparing two models of HGT rates: one as a function of genome size and ecosystem gain, and another with genome size alone. Tests were performed separately for Clusters of Orthologous Groups (COG) functional categories. Asterisks (*) denote categories where ecosystem gain significantly influenced HGT rates (*p <* 0.05 after FDR correction via the Benjamini-Hochberg method). Notably, transitions to new ecosystems were associated with lower median HGT rates in all of the categories shown here.

## Discussion

Bacterial adaptation to environmental transitions represents a fundamental process in microbial evolution and ecology. Previous studies examining the impact of environmental changes on genome dynamics have typically focused on single species or specific classes of genes. To systematically investigate this phenomenon, we analyzed horizontal gene acquisitions of diverse bacterial lineages along a deep phylogeny, testing three key expectations: (1) relaxed purifying selection and/or increased positive selection during environmental transitions; (2) a tendency of colonizing bacteria to have larger genomes; and (3) elevated HGT rates at transitions to new environments.

As expected, our analyses reveal a slight relaxation of purifying selection during ecosystem transitions, consistent with potential population bottlenecks. Such bottlenecks may allow new mutations to escape elimination quickly, contributing to higher proportions of non-synonymous substitutions compared to synonymous ones. Contrary to expectations, we found that genomes transitioning to new ecosystems tend to be small. The observed trend might reflect a trade-off between adaptability, favored by large genomes, and efficiency, favored by small genomes. Although larger genomes could ease transitions by providing broader molecular toolkits or homologous gene landing pads for HGT (Maslov et al., 2009, Taylor et al., 2024), many genes may be irrelevant to the new environment, imposing maintenance costs for redundant genetic material (Stepkowski and Legocki, 2001). Accordingly, bacteria with smaller genomes may be able to replicate faster and use resources more efficiently, traits that might be favored during colonization of new environments.

Again contradicting our expectations, we found that neither ecosystem gains nor gains of pathogenicity are associated with increased HGT rates. Instead, bacteria colonizing new ecosystems exhibit significantly reduced HGT rates – a phenomenon that is entirely attributable to their smaller genome sizes (Fig. 2, scatter plot), aligning with established relationships between genome size and HGT (Cordero and Hogeweg, 2009). After controlling for genome size, only two functional categories of genes show an effect of ecosystem gains on HGT rates: “Amino acid transport and metabolism” and “Transcription” (which includes transcription factors). For these, HGT rates are actually lower at transitions to new ecosystem.

We would like to note here two limitations of our study. First, the scopes of ‘Ecosystem Type’ labels in the GOLD database can vary substantially: the scope of a label such as ‘Fermented vegetables’ may not be comparable to that of ‘Freshwater’ aquatic environments. A broader ecosystem label may result in fewer inferred transitions, missing out on the narrower niches that may be a part of the broader ecosystem. Second, while sampling issues are not a significant concern in the gene families we analyzed, given the scale and dense taxon sampling of the dataset, the same cannot be said for the ecosystem labels. The number of genomes in the dataset for each ecosystem label is not uniform, and most genomes in GOLD are sampled in only one or a few ecosystems. On one hand, missing ecosystem labels may result in certain ecosystem gains not being inferred; on the other hand, missing labels for a subset of a given clade may also lead to the inference of a false positive ecosystem gain when in reality the clade’s ancestor was already adapted to that ecosystem. While both limitations likely add noise to our analysis, its power to detect biologically relevant signals is confirmed by our observations of (i) smaller genome sizes at ecosystem gains and (ii) a strong association between genome size and HGT rates. Moreover, both limitations are unlikely to bias our results in any systematic manner.

Given that HGT is often considered the dominant mechanism of bacteria to adapt to new environments (Pang and Lercher, 2019, Pál et al., 2005, Arnold et al., 2021), our finding that HGT rates are not affected by transitions to new ecosystems (after controlling for genome size) seems counterintuitive. Moreover, both a weakening of purifying selection and an increase of positive selection – as observed in the nucleotide substitution patterns – would be expected to be associated with higher HGT rates. Why might HGT rates be largely unaffected by transitions between environments? It is possible that a colonizing bacterium may occupy a different microniche (e.g., different metabolic preferences, spatial segregation, or competitive exclusion) than resident species, reducing physical interactions needed for HGT (Dmitrijeva et al., 2024, Polz et al., 2013, Foster and Bell, 2012). Moreover, the colonizing bacterium’s recombination machinery may not be compatible with resident mobile genetic elements (Johnson and Nolan, 2009). While these conjectures might potentially explain – at least in part – the lack of evidence for increased HGT rates at transitions to new ecosystems, more research will be needed to fully understand this surprising finding. Overall, our findings suggest that colonization events of bacteria may be less driven by adaptive benefits of HGT than previously thought.

## Methods

### Data retrieval and processing

We retrieved NOGs (gene families) from the EggNOG database v6, and the corresponding gene trees. The genome tree topology was estimated using broad-distribution NOGs (at least 95% of taxa represented in each NOG) with ASTRAL-Pro 2. Branch lengths on this topology were estimated using IQ-TREE 2 using `Q.pfam+I+R8` model on a concatenated set of multiple sequence alignments retrieved from EggNOG. The genome tree was rooted using the Minimum Ancestor Deviation method (Tria et al., 2017), which has been shown to be more accurate than other rooting methods (Wade et al., 2020).

The ‘Ecosystem Type’ labels for each genome were retrieved from the GOLD database (v9). GOLD follows a hierarchical labeling system, and the ‘Ecosystem Type’ level provides the most fine-grained and diversified set of labels such that most genomes have a label at this level, unlike, for example, the ‘Ecosystem Subtype level. We used the NCBI Datasets API to retrieve nucleotide sequences for each genome (O’Leary et al., 2024). Nucleotide sequences were used to prepare codon alignments for each NOG, translated from the amino acid multiple sequence alignments provided by EggNOG using PAL2NAL (Suyama et al., 2006).

Pathogenic and commensal *E*.*coli* and *Shigella* genomes were retrieved from NCBI RefSeq using the NCBI Datasets API. Strain genomes with at least 95% CheckM completeness and without suppressed assembly status were retained. The pathogenicity of each genome was inferred from the ‘Pathogenicity’ label in the assembly attributes. OrthoFinder was used to infer orthologous groups of genes, and corresponding gene trees were estimated using IQ-TREE 2 with the ‘JTT’ substitution model.

### Inference of HGT, Environmental transitions, and Selection intensities

HGT was inferred using Count with asymmetric Wagner parsimony. For inferences of HGT, ecosystem gains, and pathogenicity gains, the gain penalty ratio for Count was set to 1.0, which means that ecosystems or pathogenicity gains are penalized the same as losses.

We estimated the direction of selection (*ω*) and the intensification/relaxation parameter (*k*) for 3228 NOGs for which we could reliably retrieve nucleotide sequences for all taxa containing the NOG, with at least two branches were in either branch set, and for which we could successfully run RELAX. For LRT of HGT rates in functional categories, we used the COG functional categories as defined in the COG database (Tatusov et al., 2000, Galperin et al., 2021). We performed LRTs for each COG category with at least 20 NOGs in our dataset, comparing a model with HGT rate as a function of genome size and ecosystem gain to a nested model with genome size only. The p-values were corrected for multiple testing using the Benjamini-Hochberg method (Benjamini and Hochberg, 1995).

LRTs testing for the effect of ecosystem gains, pathogenicity gains, as well as the effect of ecosystem gains on specific functional categories were used to compare generalised linear models (GLMs) with and without the categorical environmental gain predictor. The GLMs were fitted using the GLM function in Python Statsmodels (Seabold and Perktold, 2010) using a log-link function.

## Code and data availability

The code used in the analysis of ecosystem transitions is available in the following Gitlab repository: https://gitlab.cs.uni-duesseldorf.de/mishra/hgt_rates_and_ecosystem_transfers For pathogenicity analysis, it is at: https://gitlab.cs.uni-duesseldorf.de/mishra/hgt_rates_and_pathogenicity_gains The data used or generated in this study can be retrieved from Zenodo: https://doi.org/10.5281/zenodo.14555205.

