## Supplementary Information for "Streamlined genomes, not horizontal gene transfer, mark bacterial transitions to unfamiliar environments"

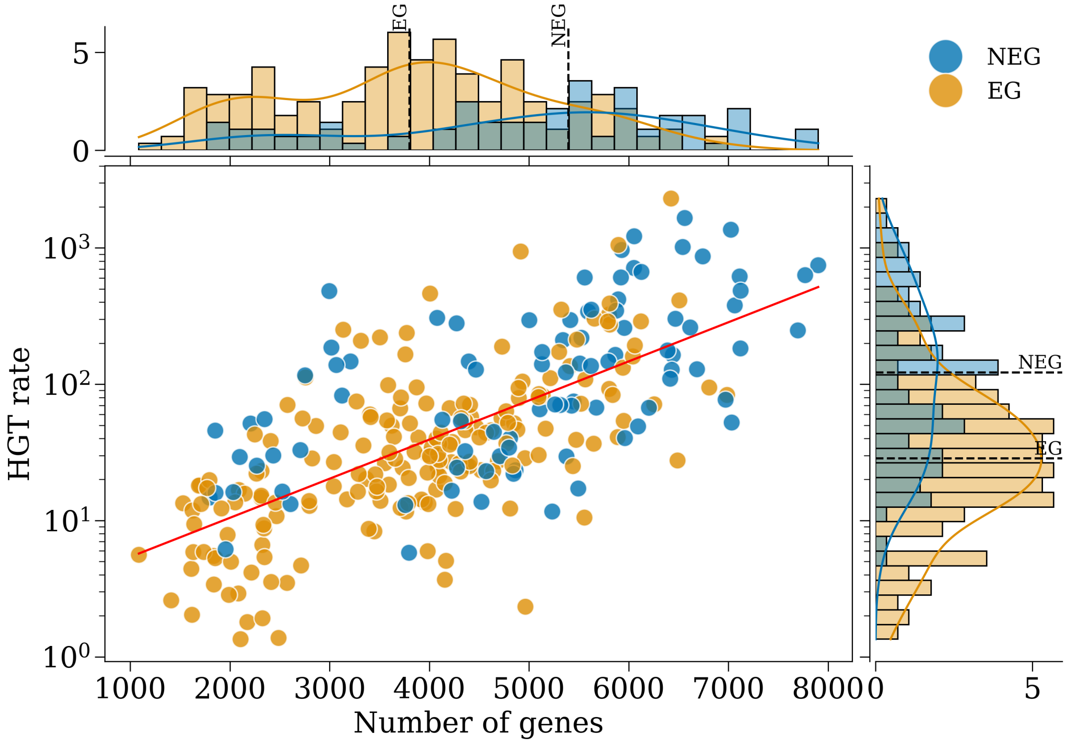


**Fig. S1. Bacteria transitioning to new ecosystems have smaller genomes and lower HGT rates (single-copy NOGs).** As seen in the top marginal histogram, branches with ecosystem gains (EG, orange) tend to have a smaller genome size than branches with no ecosystem gains (NEG, blue). Branches with ecosystem gains also tend to have lower HGT rates, as shown in the right marginal histogram (Mann-Whitney $U = 7622.5, p = 3 \times 10^{-16}, N_{EG} = 208, N_{NEG} = 108$). Dotted lines show medians. Each point in the scatter plot represents a branch in the genome tree on which at least one gene was gained through HGT. Histograms show percentages. The regression line is $log(y) = 2.79 + 6.61 \times 10^{-4} \cdot x$, with a Pearson correlation coefficient $r = 0.86$.

ALT TEXT: HGT rate is a function of genome size, as shown here in the scatter plot. Marginal plots show the distribution of genome sizes (top) and HGT rates (right) for branches with and without ecosystem gains.

| COG category | Median HGT rate NEG | EG | Log Likelihood Ratio | p-value |
|---|---|---|---|---|
| Amino acid transport and metabolism | 16.015 | 5.157 | 23.094 | 0.000 |
| Transcription | 26.941 | 6.079 | 7.995 | 0.047 |
| Carbohydrate transport and metabolism | 15.752 | 8.683 | 5.461 | 0.097 |
| RNA processing and modification | 11.416 | 3.676 | 5.706 | 0.097 |
| Posttranslational modification, protein turnover, chaperones | 16.465 | 5.469 | 4.528 | 0.133 |
| Lipid transport and metabolism | 9.843 | 6.018 | 3.828 | 0.168 |
| Defense mechanisms | 17.871 | 6.015 | 3.520 | 0.173 |
| Energy production and conversion | 13.754 | 7.943 | 2.690 | 0.202 |
| Cell motility | 14.447 | 5.617 | 2.706 | 0.202 |
| Coenzyme transport and metabolism | 13.020 | 7.511 | 2.720 | 0.202 |
| Translation, ribosomal structure and biogenesis | 12.827 | 5.809 | 1.590 | 0.377 |
| Nucleotide transport and metabolism | 6.363 | 5.742 | 1.176 | 0.428 |
| Intracellular trafficking, secretion, and vesicular transport | 7.611 | 5.134 | 1.238 | 0.428 |
| Signal transduction mechanisms | 17.261 | 10.120 | 0.977 | 0.461 |
| Cell wall/membrane/envelope biogenesis | 16.272 | 8.612 | 0.717 | 0.496 |
| Replication, recombination and repair | 12.706 | 6.999 | 0.789 | 0.496 |
| Secondary metabolites biosynthesis, transport and catabolism | 15.278 | 5.731 | 0.587 | 0.522 |
| Extracellular structures | 12.441 | 6.752 | 0.315 | 0.605 |
| Cell cycle control, cell division, chromosome partitioning | 16.842 | 8.051 | 0.325 | 0.605 |
| Inorganic ion transport and metabolism | 16.465 | 7.162 | 0.062 | 0.804 |

**Table S1.** COG categories with effect of ecosystem gain on HGT rates. All categories shown here have a lower median HGT rate at ecosystem gains (EG) than without ecosystem gains (NEG). The LRT p-values are corrected for multiple testing using the Benjamini-Hochberg method. They suggest that only the categories ‘Amino acid transport and metabolism’ and ‘Transcription’ are significantly affected by ecosystem gains.
